# Dating and quantifying past gene flow due to hybridization events in Iberian chubs

**DOI:** 10.1101/2025.04.13.648569

**Authors:** Estêvão Faustino, Sofia L. Mendes, João Carvalho, Silvia Perea, Ignacio Doadrio, Philine G. D. Feulner, Carla Sousa Santos, Vitor C. Sousa

## Abstract

Hybridization - the successful reproductive cross between individuals from different species - is increasingly recognized as a relatively common phenomenon in nature, mostly due to the increasing availability of genomic data for many organisms and the development of sophisticated methods to detect past gene flow. As a result, evidence of past gene flow has been detected in the genomes of many species, allowing us to investigate the evolutionary forces that act upon hybrid genomes. Biological systems where multiple instances of hybridization have been detected at different time scales are especially useful for this, as they allow investigating the effects of selection and recombination at different time scales post-hybridization. The Iberian chubs (*Squalius spp.*) present an ideal study system for this. In this group of primary fish inhabiting the rivers of the Iberian Peninsula, multiple instances of past hybridization at different geographical and time scales were reported between different species. However, before we investigate the effects of selection and recombination, it is crucial that we date the hybridization events and quantify the contribution from each parental species. In this study, we infer the demographic history of two Iberian chub hybrid lineages, one resulting from a proposed ancient hybridization event (*S. pyrenaicus*) and another potentially much more recent (São Martinho). We generated whole genome resequencing data for 52 individuals from the two hybrid lineages and their putative parental populations. Our demographic modelling results confirm the hypothesis of ancient hybridization between *S. carolitertii* and *S. tartessicus* around 600,000-800,000 years ago, giving rise to the *S. pyrenaicus* lineage. We also confirm that *S. carolitertii* is the major parental, with *S. tartessicus* only contributing <10% to the *S. pyrenaicus* genome. Furthermore, we confirm the more recent hybridization between *S. caetobrigu*s and *S. pyrenaicus* (around 4500 years ago) in the São Martinho population. We observe similar contributions from both parental species (around 50%) to the genome of the individuals of this population. Overall, our results give us a deeper understanding of how and when these past gene flow events happened and the contributions of each parental species to the genomes of the individuals of the hybrid lineages.

## Introduction

The discussion regarding the relevance of hybridization (i.e., crosses between species that lead to gene flow) as an important driver of diversity is not new (Anderson & Stebbins, 1954; Stebbins, 1959). However, there has been a newfound interest in this phenomenon in recent years, mostly due to the growing availability of population genomics data for non-model organisms and development of sophisticated bioinformatic methods that allowed to detect gene flow. As a result, a large body of work has been dedicated to detecting hybridization across multiple taxa with different levels of divergence (reviewed in Taylor & Larson, 2019). From this evidence, we now know that hybridization is relatively common in the wild, and its consequences to evolutionary processes such as fueling rapid diversification (e.g., adaptive radiations) (e.g. Meier, Marques, et al., 2017; Svardal et al., 2020) and facilitating adaptation (e.g. Jones et al., 2018; Norris et al., 2015; Oziolor et al., 2019) have been described in various systems.

The study of hybridization is now moving from mostly reporting findings of hybridization towards disentangling the evolutionary forces that act upon hybrid genomes. Recombination and selection are key determinants that shape the genomic patterns of diversity, differentiation and ancestry after hybridization (Moran et al., 2021). The genome of a first-generation hybrid individual is comprised of segments from both parental species. Initially, in a first-generation hybrid, the ancestry of these segments is balanced, with 50% contribution from each parental species. Over time, recombination will break down large ancestry tracts into smaller fragments. Furthermore, depending on which species the initial hybrid(s) mate with, the genome of the individuals of the next generations might be more skewed towards one or the other initial parental species. For instance, preferential backcrosses with one of the parental species are expected to increase the contribution of that species to the admixed genomes, thus becoming the dominant (major) parental in terms of the amount of ancestry tracks. Yet, alleles from the minor parental species might become fixed due to drift. Thus, the genomes of admixed individuals are a mosaic of ancestry tracts from each parental species.

The distribution (size and number) of these ancestry tracts along the genome provides information about the demographic and selective history acting on admixed genomes. Selection can either remove deleterious variation (mostly from the minor parental) and/or fix certain beneficial mutations in the hybrid lineage. Patterns arising from the action of selection are expected to depend on recombination rate (e.g., lower contribution of minor parental in low recombining regions), whereas stochastic neutral processes (e.g. drift) are not expected to lead to correlations with recombination rate. To obtain null neutral expectations of the distribution of the number and length of ancestry tracts from each parental species along the genome, it is crucial to infer the demographic history of populations, namely to date the hybridization events and quantify the contribution from each parental species. There are now sophisticated methods to reconstruct the demographic history based on population genomics single nucleotide polymorphism (SNP) data (e.g., dadi, fastsimcoal2), accounting for gene flow between species.

Some of the most powerful biological systems to understand the evolutionary forces acting after hybridization are those where multiple instances of hybridization have occurred at different time scales. Such biological systems allow investigating the effects of selection and recombination at different time scales post-hybridization, as well as assessing the repeatability of patterns across different hybridization events. Such systems with multiple instances of hybridization have been found in nature. For example, multiple instances of hybridization between different species or even hybrid lineages with different levels of divergence and at different time scales have been described in *Lycaeides* butterflies (Chaturvedi et al., 2020), sunflowers (Owens et al., 2023) and Xiphophorus swordtail fishes (Banerjee et al., 2023; Cui et al., 2013; Langdon et al., 2022).

The Iberian chubs (*Squalius spp*) are a group of small to medium-sized primary fish endemic to the river systems of the Iberian Peninsula, in the southwestern tip of Europe. This group comprises several species with non-overlapping distributions – that is, each river basin is only inhabited by one Iberian chub species. These species are closely related to each other, forming a monophyletic group, with the latest estimates dating the most recent common ancestor of all the species to 14 MYA based on concatenated matrixes of 6 or 7 nuclear genes and the mitochondrial cytochrome b gene (MT-CYB) (Sousa-Santos et al., 2019; Perea et al., 2021). Within this monophyletic group, there are two lineages: one formed by *S. aradensis* and *S. torgalensis*, which are confined to rivers in the southwesternmost tip of Iberia (Figure 1), and another lineage comprising all the remaining Iberian chub species.

**Figure 1.**
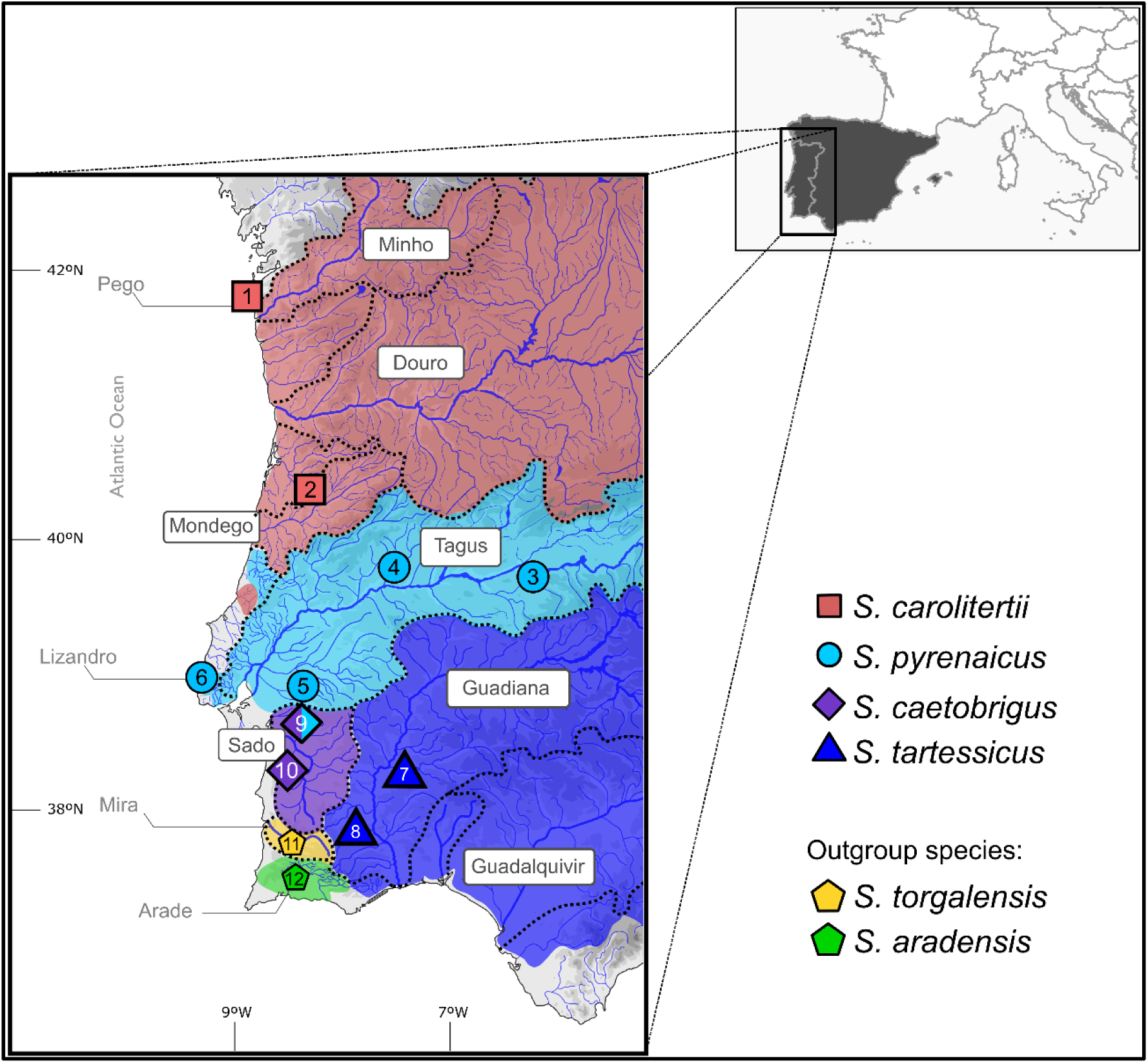
– Iberian chub species sampled for this study and respective sampling locations. Each symbol represents a sampling location, and the color and shape of the symbols represent the species sampled at that location. S. carolitertii was sampled in the (1) Pego and (2) Mortágua rivers. *S. pyrenaicus* was sampled in the (3) Ibor, (4) Pônsul, (5) Canha and (6) Lizandro rivers. *S. tartessicus* was sampled in the (7) Ardila river and (8) Oeiras stream. Finally, *S. caetobrigus* was sampled in (9) São Martinho and (10) Grândola streams. The outgroup species *S. torgalensis* and *S. aradensis* were sampled in the (11) Mira and (12) Arade River basins respectively. Sample IDs, source (newly sampled, previous studies or MNCN), sampling permits and GPS coordinates are in Table S1.

Despite being primary fish with allopatric distributions, multiple instances of hybridization have been described in the group. Mendes et al., 2021 showed that one of the species, *S. pyrenaicus*, which currently inhabits a broad area within central Iberia (Figure 1), is most likely the result of ancient admixture between two other Iberian chubs’ species – *S. carolitertii* and *S. tartessicus*. Combining genotyping by sequencing (GBS) data and demographic modelling, the authors found evidence for a contribution of both *S. carolitertii* and *S. tartessicus* to the genome of *S. pyrenaicus*. The authors also found this contribution to be imbalanced, with a higher contribution from *S. carolitertii* (∼80-90% - major parental) and a much lower contribution from *S. tartessicus* (∼20-10% - minor parental). However, their study was limited by the reduced number of SNP markers due to the data being GBS and by the absence of a reference genome at the time. Thus, it lacked the power to distinguish under which scenario the hybridization might have happened, that is, if *S. carolitertii* and *S. tartessicus* hybridized giving rise to the *S. pyrenaicus* lineage or if *S. pyrenaicus* first diverged from one of the two parental species and later upon secondary contact received a contribution from the other species. Furthermore, this study also had a restricted sampling of the putative parental species, especially of *S. carolitertii*, since only one population from the Mondego River Basin was available, which is at the southern edge of the *S. carolitertii* distribution and the northern edge of the *S. pyrenaicus* range. Since high levels of genetic differentiation between the Mondego River basin *S. carolitertii* populations and the remaining *S. carolitertii* populations have been reported based on mitochondrial and nuclear genes (Almada & Sousa-Santos, 2010; Perea et al., 2021), whether the observed evidence of hybridization is supported with alternative *S. carolitertii* populations was unclear.

A more recent study used low coverage whole genome resequencing data to investigate the prevalence of hybridization across the majority of the Iberian chub species (Mendes et al., in review). Using D-statistics, the authors found consistent evidence of *S. carolitertii* and *S. tartessicus* contribution to the genome of the *S. pyrenaicus* individuals from multiple locations across the whole distribution area of the species, which is congruent with the findings of Mendes et al., 2021 based on GBS data. As the pattern observed was consistent between *S. pyrenaicus* individuals from independent river basins, the authors propose that the hybridization event was more likely in the ancestral population, predating the range expansion and dispersal of the species its current distribution. However, exactly when and how the hybridization event happened remains unknown.

Furthermore, this same study found evidence of multiple other cases of hybridization in the Iberian chubs’ system. Specifically, the authors found evidence for recent, possibly ongoing, hybridization between *S. pyrenaicus* and *S. caetobrigus* restricted to a localized stream (São Martinho). Individuals from this population showed similar contributions from both *S. pyrenaicus* and *S. caetobrigus* into their genomes and both mitochondrial haplotypes (*S. pyrenaicus* and *S. caetobrigus* mtDNA) were found in the population at similar frequencies. Therefore, the putative ancient hybrid lineage of *S. pyrenaicus* appears to have hybridized with *S. caetobrigus*. Although this is expected to be more recent, the timing and mode of this hybridization remains unknown. Thus, the fact that multiple cases of hybridization spanning different time points and geographical ranges have been reported in the Iberian chubs makes them an ideal system to study the consequences and repeatability of hybridization. However, to investigate the consequences of hybridization, it is fundamental that we first understand the timing and mode of origin of the putative hybrid lineages.

Demographic modelling provides a powerful tool to investigate the origin of hybrid lineages. Modelling scenarios of divergence with and without gene flow allowed, for example, to investigate the role of gene flow in the ecological speciation of sticklebacks (Marques et al., 2019) and in the parallel origin of cichlids species pairs (Meier, Sousa, et al., 2017), as well as introgression with a now extinct ghost lineage in the intertidal goby *Chaenogobius annularis* (Kato et al., 2024). These methods have also been previously employed to investigate the divergence process in the Iberian chubs system, albeit with limited success due to the limited number of genetic markers (Mendes et al., 2021).

The main goal of this study was to investigate the timing and mode of the *S. pyrenaicus* and the São Martinho hybridization events. To do so, we generated high coverage whole genome resequencing data of individuals representative of the putative parental and hybrid populations involved in both cases. Using this data, we first characterized the overall patterns of population structure, genetic diversity and differentiation. We then explicitly tested for introgression in the two previously described cases of hybridization. Finally, we used demographic modelling to compare alternative divergence scenarios for each of the hybridization cases and estimate the timing and mode of hybridization (i.e., hybrid origin vs secondary contact) as well as the contribution of each parental species.

## Methods

### Sampling

Our dataset consists of 52 individuals in total. We mostly sampled 5 individuals per sampling location, with a few exceptions (see below). All samples consisted of ethanol-preserved dorsal fin clips, and we combined newly collected fin clips with samples previously collected and/or sequenced in other projects, as well as samples stored at the Museo Nacional de Ciencias Naturales (MNCN) in Madrid. To obtain the fin clips, fish were sampled using electrofishing with low-duration pulses and a small piece of the dorsal fin was collected. Individual sample IDs, source (sampled for this study, previous studies or museum), GPS coordinates of the sampling locations and permits from the Portuguese authority for species conservation (Instituto de Conservação da Natureza e das Florestas – ICNF) can be found in Table S1.

To investigate the *S. pyrenaicus* hybridization, we obtained samples representative of the distribution of *S. pyrenaicus* (Ibor – n=2, Ponsul – n=5, Canha – n=5, Lizandro – n=5), as well as the two putative parentals: *S. carolitertii* (Pego – n=5, Mortágua – n=5) and *S. tartessicus* (Ardila – n=2, Oeiras – n=5). To investigate the São Martinho hybridization, we obtained samples from the São Martinho population (n=10) and one of the putative parentals - *S. caetobrigus* (Grândola – n=5). For the other putative parental, we re-used *S. pyrenaicus* Canha (n=5). Finally, we also included two outgroup species – *S. torgalensis* (n=2) and *S. aradensis* (n=1).

### DNA extraction and whole genome sequencing

We extracted total genomic DNA from the ethanol-preserved fin clips using the Qiagen DNeasy® Blood & Tissue and Zymo Quick-DNA™ Miniprep Plus commercial kits following the manufacturer’s instructions. We accessed the DNA fragmentation on a 1% agarose gel and the DNA contamination on a NanoDrop™ 4000 spectrophotometer (Thermo Fisher Scientific). We quantified the DNA concentration on a Qubit® 2.0 Fluorometer (Thermo Fisher Scientific) using the broad-range kit assay.

The library preparation and whole genome sequencing were performed at Novogene, UK. To construct the libraries, the DNA was randomly sheared, the resulting fragments were end-repaired and A-tailed and ligated with the Illumina adapters. The fragments with Illumina adapters were then size-selected, PCR amplified and purified. The resulting library was paired-end sequenced at 150 base pairs (bp) on Illumina NovaSeq6000.

### Mapping and filtering

We inspected the quality of the sequencing data using fastqc v0.11.9 (https://www.bioinformatics.babraham.ac.uk/projects/fastqc/). We then used fastp v0.20.1 (Chen et al., 2018) to remove reads with 40% or more of the bases with Phred-score below 20 and to trim poly-G tails.

We used the genome of *Squalius cephalus* (GenBank assembly accession GCA_022829025.1), the central European chub, as a reference to map our reads. To mask regions of low complexity or repetitive regions of the genome, as these can affect demographic inferences (Patil & Vijay, 2021), we used *RepeatMasker* v.4.1.5 (https://repeatmasker.org) on the reference genome of *S. cephalus*. A total of 247,891,138 bp, representing 23.84% of the genome, was masked for repetitive regions such as retroelements and DNA transposons.

We then aligned the reads from each individual to the *S. cephalus* genome using BWA-MEM from BWA v0.7.17 (Li, 2013) with default parameters and sorted the output alignments using samtools v1.12 (Li et al., 2009). For some individuals, the library had previously been sequenced at lower coverage for a different project, stored, and now re-sequenced at higher coverage. This produced multiple sets of reads. In such cases, we mapped them separately and then merged the coordinate-sorted alignment using samtools v1.12 (Li et al., 2009). For each individual alignment, we then verified and fixed mate-pair information with FixMateInformation, sorted the output by query name, and marked and removed duplicated reads with MarkDuplicates from Picard v3.1.1 (https://broadinstitute.github.io/picard/). We also evaluated the quality of the resulting filtered coordinate sorted alignment using Picard v3.1.1 (CollectWgsMetricsWithNonZeroCoverage, CollectAlignmentSummaryMetrics). Finally, we indexed the final alignment of each individual using samtools v1.12 (Li et al., 2009).

Before calling genotypes, we used samtools v1.12 (Li et al., 2009) to remove reads that mapped in the repetitive regions identified by RepeatMasker, coordinate sort and index the output alignments. These individual alignments were then used for genotype calling. We used freebayes v1.3.6 (Garrison & Marth, 2012) to perform individual haplotype-based genotype calling requiring a minimum mapping quality of 30, a minimum base quality of 20, including monomorphic sites and without any Hardy– Weinberg equilibrium priors (-p 2 --min-mapping-quality 30 --min-base-quality 20 -- report-monomorphic --hwe-priors-off). We performed the genotype calling to create two different datasets, which were filtered separately. For the *S. pyrenaicus* dataset, we retained all the *S. carolitertii*, *S. pyrenaicus* and *S. tartessicus* individuals, as well as the outgroup individuals. For the São Martinho dataset, we retained the *S. caetobrigus*, *S. pyrenaicus* Canha and São Martinho individuals, as well as the outgroups.

We first filtered the output VCF files to keep only biallelic SNPs and monomorphic sites. Then, using vcftools v.0.1.16 (Danecek et al., 2011), we filtered each individual based on their mean coverage, filtering out sites below a minimum coverage threshold of 10x, and a maximum coverage of twice its mean coverage. Next, to remove sites with excessive heterozygosity across the entire dataset that are likely due to mapping errors, we performed a test to detect deviations (excess of heterozygotes) from Hardy-Weinberg (HW) proportions implemented in vcftools in the pooled samples. We used vcftools to ensure sites in repetitive regions were discarded. We applied a hard filter of a maximum of 25% missing data allowed per site. After this, we recorded the number of monomorphic sites that passed all these filters and kept only SNPs. Since the HW test was conducted with all pooled samples, this step was repeated by sampling locations for the *S. pyrenaicus* dataset. For this, we merged sampling locations with two or fewer individuals with the closest sampling location (Ibor and Ponsul originated the “Mid Tagus” sampling location and Ardila and Oeiras gave rise to the “Guadiana” sampling location). Then, we removed sites that are heterozygote at all individuals in any two populations and may be due to mapping errors. This resulted in the removal of 10,926 sites using vcftools. This second HW test was not applied to the São Martinho dataset. Since we expect this to be a more recent hybridization event, we expected that a higher proportion of sites would display this pattern. As such, for the São Martinho dataset we decided not to remove sites with an excess of heterozygotes when pooling pairs of populations.

We used the two closer outgroup species (*S. torgalensis* and *S. aradensis*) to polarize the ancestral and derived alleles at each SNP. Since *S. aradensis* and *S. torgalensis* have diverged more recently (∼14 Mya) from our focal species than *S. cephalus,* used as the reference genome (∼19 Mya) (Sousa-Santos et al., 2019), we minimized the chance of having wrong polarization by using as ancestral the allele fixed in *S. aradensis* and *S. torgalensis* rather than using the reference allele. In more detail, we only kept sites where all individuals in the outgroup were homozygous, considering that allele as the ancestral. We discarded heterozygous sites in the outgroup, avoiding sites with higher mutation rates and possible back mutations. To keep only biallelic SNPs we filtered out sites that became monomorphic after all the filters using a custom R script. Finally, to discard wrongly called heterozygote genotypes with high allelic imbalance, we filtered sites based on a disproportionate allelic depth, setting as missing data genotypes that presented an imbalance greater than 80:20, using custom R scripts. After the filtering, the *S. pyrenaicus* dataset comprised 8,398,720 SNPs and 37 individuals and the São Martinho dataset comprised 23 individuals and 5,702,205 SNPs.

To investigate the global patterns of population structure, we created a third dataset combining the SNPs in common between the *S. pyrenaicus* (*S. pyrenaicus* dataset) and São Martinho (São Martinho dataset) datasets. We used a combination of bcftools isec and bcftools merge (Danecek et al., 2021) to obtain the SNPs from each dataset that were present in both datasets. We then performed the polarization step again on this new set of SNPs, followed by setting as missing data the heterozygotic genotypes with an allelic imbalance greater than 80:20. This resulted in the Merged dataset, which comprised 52 individuals and 2,399,889 SNPs.

### Overall patterns of genetic diversity and differentiation

We assessed the overall population structure through individual-based methods using the Merged dataset. Given that the methods we used to detect population structure rely on the assumption of independence of SNPs, we discarded the outgroup individuals (fixed homozygote for the ancestral allele) and performed a thinning by selecting 1 in every 100 SNPs and applied a Minor Allele Frequency filter (MAF)>5%, resulting in 18,179 SNPs. We then performed a Principal Components Analysis (PCA) using the R package LEA (Frichot & François, 2015). We also estimated population structure for 2<K<14 using ADMIXTURE v1.3.0 (Alexander et al., 2009), performing 100 independent runs for each K value. We then selected the run with the lowest cross-validation error for each K value.

To verify that the *S. pyrenaicus* and São Martinho datasets showed similar population structure patterns as the Merged dataset, we analyzed the *S. pyrenaicus* dataset and the São Martinho dataset, with the same methods following the same approach, i.e., removing the outgroup individuals, thin down to 1 in every 100 SNPs and MAF>5% resulting in 48,997 SNPs and 30,816 SNPs, respectively. For ADMIXTURE we tested 2<K<14 for the *S. pyrenaicus* dataset and 2<K<6 for the São Martinho dataset.

To quantify the levels of genetic differentiation, we calculated pairwise F_ST_ between all pairs of sampling locations for the *S. pyrenaicus* and São Martinho datasets using the Hudson estimator (Bhatia et al., 2013; Hudson et al., 1992). To quantify the levels of genetic diversity, we calculated expected and observed heterozygosity for the same two datasets using custom R scripts. To do so, we obtained the heterozygosities based on the SNPs datasets, and normalized it by the ratio of SNPs to total sites that passed our filters. This way, we obtained the ratio of heterozygote positions across the total number of sites.

### Detection of gene flow

Previous work identified introgression in *S. pyrenaicus* and in São Martinho (Mendes et al., 2021; Mendes et al., in review). To confirm whether these patterns of gene flow were also recovered in our data, we performed D-statistic (ABBA-BABA) tests (Durand et al., 2011). This test requires a user-defined bifurcating topology involving three ingroup populations (P1, P2 and P3) and an outgroup (O), related between themselves as follows: (((P1,P2),P3),O). If this is the correct topology, without gene flow, the allele sharing between the ingroup populations and P3 (i.e. sites with pattern ABBA and BABA) will be due to incomplete lineage sorting (ancestral polymorphism) and D will be zero. However, if there is excessive allele sharing between P1 and P3, D will be negative. Likewise, if there is excessive allele sharing between P2 and P3, D will be positive. Assuming the infinite sites model, deviations of the D-statistic from zero support gene flow either between P3 and P1 (D<0) or P3 and P2 (D>0). In our tests, our focal population is always P2 and thus an excess of allele sharing due to gene flow will result in positive D values. Furthermore, all calculations were performed using *S. torgalensis* as the outgroup species. We assessed the significance of all the D-statistic values using a block jackknife approach (Soraggi et al., 2018) with a 5 Mb block size.

To test for evidence of gene flow involving *S. pyrenaicus*, we tested *S. pyrenaicus* as P2, while testing both *S. carolitertii* and *S. tartessicus* as P1 and P3, using the *S. pyrenaicus* dataset. We tested all possible combinations of sampling locations from these three species to assess the effects of the sampling locations used to perform the test.

To test for evidence of gene flow involving the São Martinho individuals, we tested São Martinho as P2, while testing both *S. caetobrigus* and *S. pyrenaicus* Canha as P1 and P3 using the São Martinho dataset.

For both hybridization cases, we also calculated the D-statistic per individual to assess if there was any individual variation. Likewise, we also estimated the contribution of each parental species to the genome of the putative hybrid individuals. For *S. pyrenaicus* (*S. pyrenaicus* dataset), we estimated the contribution of *S. tartessicus* (*α*) to the genome of each *S. pyrenaicus* individual using the f4-ratio between the difference of ABBA and BABA sites in the topology (((*S. carolitertii*, *S. pyrenaicus*), *S. tartessicus*), Outgroup) and the same difference under complete admixture (Patterson et al., 2012). To estimate complete admixture, we split the *S. tartessicus* Oeiras individuals into two groups (Oeiras 1 (2 individuals) and Oeiras 2 (3 individuals)) and calculated the difference between ABBA and BABA sites for a topology where P2 is substituted by one group and P3 by the other, with P1 remaining *S. carolitertii*: (((*S. carolitertii*, *S. tartessicus 1*), *S. tartessicus* 2), Outgroup). The f4-ratio between the difference of ABBA and BABA sites in these two topologies gives us an estimation of the *S. tartessicus* contribution. For the São Martinho hybridization case (São Martinho dataset), we estimated the contribution of *S. pyrenaicus* (*α*) to the genome of each São Martinho individual in a similar manner.

### Demographic modelling

For both hybridization cases, we compared alternative divergence scenarios between three populations through demographic modelling based on the site frequency spectrum using *fastsimcoal2* v2.8 (Excoffier et al., 2013, 2021). To generate the three-population Site Frequency Spectrum (3D-SFS), we performed a block resampling approach, dividing the genome into 100kb continuous blocks. Assuming block independence, we randomly selected individuals from each population in each block, keeping only SNPs from blocks with a median distance between SNPs greater than 2 bp. We downsampled to a lower number of individuals (see below), keeping only SNPs without missing data, thus resulting in a 3D-SFS without missing data. For each case (*S. pyrenaicus* dataset or São Martinho dataset), we then added the corresponding number of monomorphic sites previously calculated to the obtained 3D-SFS.

We then compared different models. For each model, we performed 100 independent runs with 50 cycles, out of which the first 20 cycles considered monomorphic sites when computing the composite likelihood, whereas for the remaining 30 cycles only SNPs were considered to compute the composite likelihood, approximating the expected SFS with 100,000 coalescent simulations. We assumed a mutation rate of 1e-9 mutations per site per generation, according to estimates for other fish species (Bergeron et al., 2023; Feng et al., 2017; Malinsky et al., 2018; Zhang et al., 2023). To convert the divergence times into absolute time in Mya, we assumed a generation time of three years (Almada & Sousa-Santos, 2010).

For the *S. pyrenaicus* hybridization case, we compared 8 alternative divergence scenarios between *S. carolitertii*, *S. pyrenaicus* and *S. tartessicus.* Using the dataset *S. pyrenaicus*, we selected the Pego sampling location of *S. carolitertii*, the Lizandro sampling location for S*. pyrenaicus* and the Oeiras sampling location for *S. tartessicus* and generated the 3D-SFS randomly sampling 4 individuals from each population per block. This resulted in an observed SFS with 2,118,081 SNPs and 417,634,738 monomorphic sites. We first tested two models without any gene flow, where *S. carolitertii* and *S. tartessicus* diverge from each other and *S. pyrenaicus* then diverges from either *S. carolitertii* (model ((Carol,Pyr),Tar)) or *S. tartessicus* (model (Carol,(Pyr,Tar))). We then tested three models with gene flow. We compared a model where *S. pyrenaicus* receives a contribution *α* from *S. tartessicus* and 1-*α* from *S. carolitertii* at the time of hybridization (model Hybrid Origin) to two other models where *S. pyrenaicus* first diverges from *S. carolitertii* (model ((Carol,Pyr),Tar)+SecContTar) or *S. tartessicus* (model (Carol,(Pyr,Tar))+SecContCarol) and later receives a contribution *α* from the other species (secondary contact models). Finally, we tested three models identical to the three models with gene flow but allowing for divergence with migration between *S. carolitertii* and *S. tartessicus* until the time of the split of *S. pyrenaicus*. All eight models include a recent bottleneck in the *S. carolitertii* Pego population. This is because Pego is a small, independent coastal river (less than 8km long) and, therefore, has a low carrying capacity. Furthermore, all eight models allow for population size resizes of the ancestral *S. carolitertii* and *S. tartessicus* populations during population splits in the secondary contact models and the models without admixture. The Hybrid Origin models allowed resizes during the admixture pulses.

For the São Martinho hybridization case, we compared 6 alternative divergence scenarios between *S. caetobrigus*, *S. pyrenaicus* and the São Martinho population. Using the “São Martinho” dataset, we randomly sampled 4 individuals from *S. caetobrigus* and *S. pyrenaicus* and 8 individuals from São Martinho per block to generate the 3D-SFS. This resulted in an observed SFS with 846,554 SNPs and 418,257,746 monomorphic sites. We first compared three models. We compared a hybrid origin model where São Martinho receives a contribution *α* from *S. pyrenaicus* and 1-*α* from *S. caetobrigus* at the time of hybridization (model Hybrid Origin) to two other models where the São Martinho population first diverges from *S. caetobrigus* (model ((Caet,SãoMartinho),Pyr)+SecContPyr) or *S. pyrenaicus* (model (Caet,(SãoMartinho,Pyr))+SecContCaet) and later receives a contribution *α* from the other parental population (secondary contact models). We then compared three other models, in all similar to the previous ones but allowing migration between *S. caetobrigus* and *S. pyrenaicus* until the split of the São Martinho population.

## Results

### Overall patterns of genetic diversity and differentiation

The overall population structure (Merged dataset) revealed intra-specific population structure within *S. pyrenaicus* (Figure 2A). The best K value (K=6, Figure S3) separated *S. carolitertii*, *S. caetobrigus* and *S. tartessicus*, as well as three clusters within *S. pyrenaicus*, corresponding to the Middle Tagus (Ibor + Ponsul sampling locations), Lower Tagus (Canha sampling location) and Lizandro River basin (Figure 2A). This separation is also supported by higher K values (Figure S4). The São Martinho individuals showed evidence of being admixed, with almost equal contributions from *S. caetobrigus* and *S. pyrenaicus* Canha, which varied between individuals. The assignment of Canha and these proportions of the São Martinho individuals to the same cluster supports our choice of using the Canha sampling location as the representative population of the *S. pyrenaicus* parental for the São Martinho hybridization case.

**Figure 2.**
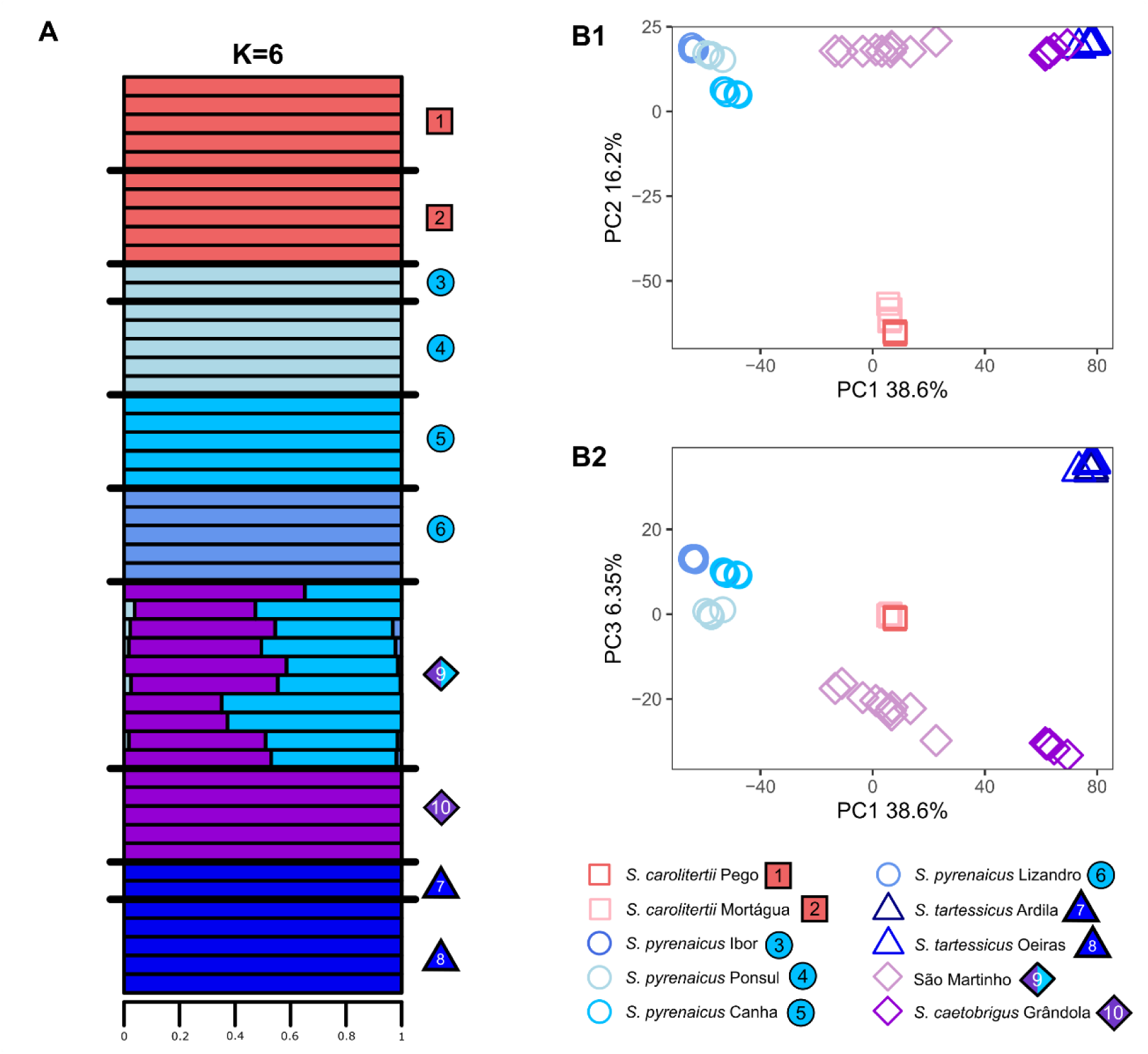
– Overall population structure inferred from the merged dataset (dataset Merged). (A) Admixture plot obtained with ADMXITURE for K=6. Each horizontal bar corresponds to one individual. Individuals are ordered from north to south. (B) Principal Components Analysis with all focal species analyzed. Each symbol corresponds to one individual. B1) PC1 and PC2; B2) PC1 and PC3.

The PCA based on Merged dataset also supported the same overall pattern, with PC1, PC2 and PC3 providing separation between the four species (Figures 2B and S4) and latter PCs providing separation between *S. pyrenaicus* sampling locations (PC4 and PC5 – Figure S5). The admixed status of the São Martinho individuals is also reflected on the PCA, with these individuals appearing in between the *S. pyrenaicus* and *S. caetobrigus* clusters.

The patterns of population structure recovered from the *S. pyrenaicus* and São Martinho datasets are overall congruent with those of the Merged dataset (Figures S6 to S11). For the *S. pyrenaicus* dataset, we also observe separation between the different sampling locations of *S. pyrenaicus*. This is supported by the PCA as well as by the K values with lower cross-validation errors (Figures S6-S8). Furthermore, for this dataset, we also observed separation between the Pego and Mortágua sampling locations of *S. carolitertii* at lower K values (Figure S8). These two sampling locations are also separated by PC3 (Figure S6). This is not surprising, as these two sampling locations are independent river basins located the edges of the *S. carolitertii* distribution. Furthermore, this pattern of high genetic differentiation between *S. carolitertii* from tributaries of the Mondego River basin, as is the case of Mortágua, as the remaining *S. carolitertii* had already been reported (Perea et al., 2021). Regarding the São Martinho dataset, the analyses also support the clear separation of the two parental species and the admixed status of the São Martinho individuals is also recovered (Figures S9-S11).

We observe overall high levels of genetic differentiation (Table S3), recovering the relative results previously reported by Mendes et al using low coverage whole genome data (Mendes et al., in review). For the *S. pyrenaicus* dataset, the lowest levels of genetic differentiation were found within species, in particular between the two *S. tartessicus* sampling locations (F_ST_=0.128 - Table S3A). Within *S. pyrenaicus*, the levels of genetic differentiation between sampling locations are congruent with the clusters observed in the admixture analysis (Figure S4 and S8). For example, the lowest levels of differentiation were observed between the Ibor and Ponsul sampling locations (F_ST_=0.216), which often clustered together in the admixture analysis. Regarding the São Martinho dataset, we observed substantially lower levels of genetic differentiation between the São Martinho individuals and both of its parentals (F_ST_≤0.376) than between the two parental species (F_ST_=0.781 - Table S3A). This is consistent with the admixed status of these individuals and the PCA and Admixture results (Figure 2, S4, S5, S9 and S11).

Regarding the levels of genetic diversity, for the species with broader distribution ranges (*S. carolitertii*, *S. pyrenaicus* and *S. tartessicus*) we observed an overall pattern of increasing genetic diversity from north to south, with the northernmost species, *S. carolitertii,* having the lowest levels of genetic diversity and *S. tartessicus* the highest (Table S4A). The São Martinho population was an outlier to this north-south pattern, as it showed the highest levels of genetic diversity (Table S4B), which is congruent with its admixed status (Figure 2). We also observed that *S. caetobrigus* has lower levels of genetic diversity than *S. pyrenaicus* and *S. tartessicus*, with levels of diversity only slightly higher than those of *S. carolitertii* (Table S4). However, despite its geographical distribution being close to those of *S. pyrenaicus* and *S. tartessicus*, *S. caetobrigus* occupies a much smaller distribution area, being restricted to the Sado river basin, and thus likely has a smaller effective size than *S. pyrenaicus* or *S. tartessicus*.

### Detection of gene flow

Given the previous studies suggesting that both *S. pyrenaicus* and the São Martinho population are the result of hybridization, we explicitly tested for gene flow due to hybridization in both lineages using the D-statistics.

When testing for evidence introgression on the genome of *S. pyrenaicus* using both *S. carolitertii* or *S. tartessicus* as sister taxa, we obtained consistently positive significant D values, indicating excessive allele sharing between *S. pyrenaicus* and both *S. carolitertii* and *S. tartessicus* (Figures 3A and S12). The D values obtained were not affected by the *S. pyrenaicus* population used, and we observed little to no individual variation in D values (Figure 3A, Table S5), indicating that the contribution of *S. carolitertii* and *S. tartessicus* is similar between individuals, even from independent river basins. These results are consistent with what had previously been reported by Mendes et al., in review, pointing to an ancient hybridization event.

**Figure 3.**
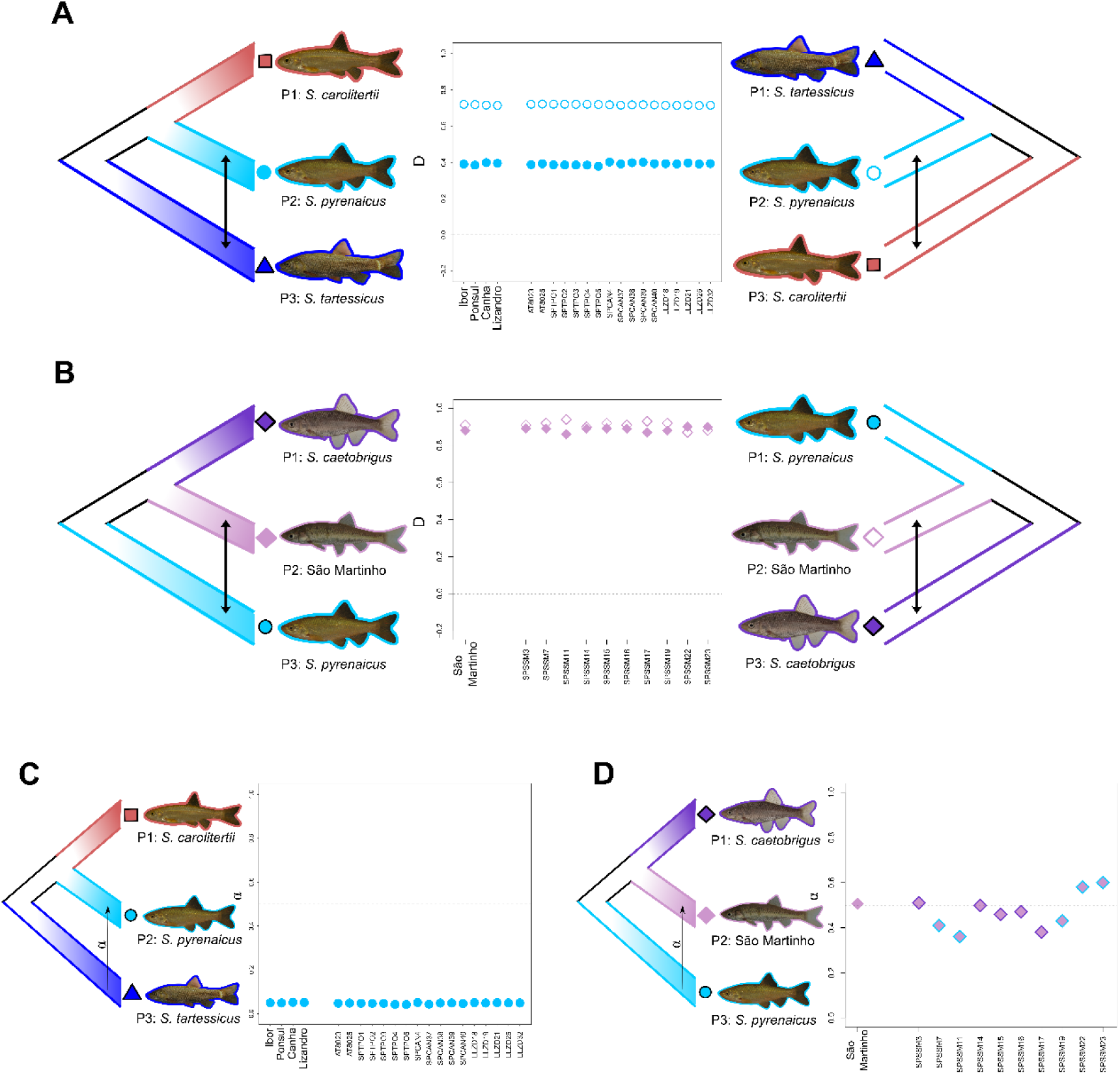
– Evidence of introgression in *S. pyrenaicus* and in São Martinho. For panels A and B, filled symbols on the plot correspond to the topology represented on the left and empty symbols correspond to the one represented on the right side of the plot. *S. torgalensis* is the outgroup used in all tests. (A) Evidence of introgression from *S. tartessicus* (filled circles) and *S. carolitertii* (empty circles) into *S. pyrenaicus*. The calculation was performed per *S. pyrenaicus* sampling location (Ibor, Ponsul, Canha and Lizandro) as well as per individual (on the x-axis). All D values are significant (p<0.01). (B) Evidence of introgression from *S. pyrenaicus* (filled diamonds) and *S. caetobrigus* (empty diamonds) into the São Martinho individuals calculated for all individuals together (São Martinho) and per individual. All D values are significant (p<0.01). (C) Contribution of *S. tartessicus* into the genome of the *S. pyrenaicus* individuals. α is the estimated proportion of *S. tartessicus* contribution to the genome of each *S. pyrenaicus* individual or group of individuals. The four *S. pyrenaicus* sampling locations and the different individuals are shown along the x-axis. The Oeiras sampling location of *S. tartessicus* and the Pego sampling location of *S. carolitertii* were used to perform all calculations. The dashed line is 0.5, which would correspond to an equal contribution from both species. (D) Contribution of *S. pyrenaicus* into the genome of the São Martinho individuals. α is the estimated proportion of *S. pyrenaicus* contribution to the genome of each São Martinho individual or the whole group of individuals. The dashed line is 0.5, which would correspond to an equal contribution from both species. The borders on the diamonds indicate the mtDNA of each individual, that is, a blue border means *S. pyrenaicus* mtDNA and a purple border means *S. caetobrigus* mtDNA.

Regarding the São Martinho individuals, our D-statistic results also corroborate their admixed status. When testing for introgression on the genome of the São Martinho individuals using *S. caetobrigus* or *S. pyrenaicus* as sister taxa, we found consistently positive significant D values indicating excessive allele sharing with both *S. pyrenaicus* and *S. caetobrigus*, respectively (Figure 3B, Table S6). However, contrary to the *S. pyrenaicus* case, we found considerable variation in parental contributions between individuals (Figure 3B), consistent with a more recent hybridization event.

When estimating the contribution of *S. tartessicus* to the genome of *S. pyrenaicus*, we found this contribution to be low and identical between individuals (∼5%) (Figures 3C, S13 and Table S7). This confirms the previous work indicating that *S. tartessicus* is the minor parental in the *S. pyrenaicus* hybridization case (Mendes et al., 2021). Contrastingly, when estimating the contribution of *S. pyrenaicus* to the genome of the São Martinho individuals, we found close to 50% average *S. pyrenaicus* contribution and considerable variation between individuals (between 36 to 60% contribution) (Figure 3D, Table S8). This shows it is difficult to assign *S. caetobrigus* or *S. pyrenaicus* into a major or minor parental category in the São Martinho hybridization case and points to a much more recent hybridization event, as the parental contributions are closer to 50% from each parental species. Furthermore, we observed no association between the proportion of contribution from each parental species and the mitochondrial type of each São Martinho individual (Figure 3D), i.e. individuals with *S. pyrenaicus* mtDNA did not have a higher contribution from that parental species.

#### Demographic modelling

In the *S. pyrenaicus* hybridization case, we started by testing models without admixture, comparing a divergence scenario with *S. carolitertii* (model ((Carol,Pyr),Tar)) or with *S. tartessicus* (model (Carol,(Pyr,Tar))). A closer relationship between *S. carolitertii* and *S. pyrenaicus* attained a higher likelihood than the alternative topology (Table S9).

The divergence of *S. pyrenaicus* from *S. tartessicus* indicated a closer relationship between *S. carolitertii* and *S. pyrenaicus*. According to this model, the ancestral of *S. carolitertii* and *S. pyrenaicus* and *S. tartessicus* diverged from each other around 2.6 MYA, with *S. pyrenaicus* diverging from *S. carolitertii* around 1.2 MYA.

We then tested two models that included a secondary contact pulse between *S. pyrenaicus* and its most distant sister species (models ((Carol,Pyr),Tar)+SecContTar and (Carol,(Pyr,Tar))+SecContCarol) and compared them to a model where *S. pyrenaicus* descends from a hybridization event between *S. tartessicus* and *S. carolitertii* (model Hybrid Origin). The three models reached similar likelihoods, placing the divergence of *S. carolitertii* and *S. tartessicus* around 2.8-2.9 MYA and estimating that *S. pyrenaicus* receives a minor contribution from *S. tartessicus* of ∼5-8%. We also noticed that the model of secondary contact between *S. carolitertii* and *S. pyrenaicus* converges with the model of hybridization, with overlapping estimates of divergence of *S. tartessicus* and *S. pyrenaicus* and the secondary contact pulse between *S. carolitertii* and *S. pyrenaicus*.

Similarly, we compared the models of secondary contact and hybridization event with continuous gene flow after the divergence of *S. carolitertii* and *S. tartessicus* until the split of *S. pyrenaicus* from one of them (models ((Carol,Pyr),Tar)+SecContTar+AncMig; (Carol,(Pyr,Tar))+SecContCarol+AncMig and Hybrid Origin+AncMig). The likelihoods of these models did not increase considerably; however, introducing ancient migration between *S. carolitertii* and *S. tartessicus* led to the three models converging to the hybridization event scenario. As such, the parameter estimates support that *S. carolitertii* and *S. tartessicus* hybridized ∼650,000 – 900,000 years ago, with a contribution of ∼6-7% from *S. tartessicus* to *S. pyrenaicus*. Most inferred parameters were consistent between models, especially times of divergence and admixture, as well as parental contribution to *S. pyrenaicus*.

Regarding the São Martinho hybridization case, we first compared a model where the São Martinho population results from a hybridization event between *S. caetobrigus* and *S. pyrenaicus* to two secondary contact models where the São Martinho population diverges from one of the parental species and later receives a contribution via secondary contact from the other parental species. We then explored three similar models but considering gene flow between the parental species *S. caetobrigus* and *S. pyrenaicus* before the time of split of the São Martinho population. Within each group of three models, the secondary contact model where the São Martinho population diverges from *S. caetobrigus* and later receives a contribution from *S. pyrenaicus* through secondary contact consistently reached a higher likelihood (Table S10). The models with gene flow between parental species reached a higher likelihood than those without gene flow (Table S10). Thus, our overall best model is the one where *S. caetobrigus* and *S. pyrenaicus* diverge from each other with gene flow approximately 1MYA, São Martinho diverges from *S. caetobrigus* ∼16,000 years ago and later receives a contribution of ∼51% from *S. pyrenaicus* around 4,500 years ago (((Caet,SãoMartinho),Pyr) + SecContPyr + AncMig). The model also infers *S. caetobrigus* always had a smaller effective size than *S. pyrenaicus*, with the ancestral effective size of *S. caetobrigus* being ∼20% of that of the ancestral effective size of *S. pyrenaicus* (Table S10). We estimated asymmetric ancient migration, with migration only from the ancestral *S. pyrenaicus* population into the ancestral of *S. caetobrigus* but not the other way around. Most parameter values inferred are remarkably similar between models, namely the contribution of each parental species to the São Martinho population, the times of divergence and the times of admixture, as well as the migration rates between the ancestral parental populations (Table S10).

## Discussion

Taken together, our population structure, D-statistics and demographic modeling results using high-coverage whole genome resequencing data analyzed in this study confirm the previous evidence of hybridization in *S. pyrenaicus* and the São Martinho populations.

Regarding *S. pyrenaicus*, the results of the D-statistics indicate a contribution of both *S. tartessicus* and *S. carolitertii* to the genome of *S. pyrenaicus* and suggest that *S. tartessicus* is the minor parental (Figure 3A). This is consistent with previous studies, where similar patterns had been observed based on genotyping by sequencing and low coverage whole genome resequencing (Mendes et al., 2021; Mendes et al., in review). However, our estimation of the *S. tartessicus* contribution based on the ABBA/BABA patterns is lower than previously reported. While the results of Mendes et al., 2021 suggest that the contribution of *S. tartessicus* to the genome of *S. pyrenaicus* was between 16% and 20%, our estimates never surpassed 5% (Figure 3C, Table S7). This could be due to the lower number of markers from more conserved genomic regions analyzed by Mendes et al., 2021. Furthermore, our results also showed remarkable consistency in the D values and estimates of *S. tartessicus* contribution across multiple *S. pyrenaicus* individuals from different sampling locations (Figures 3A,3C). This supports the hypothesis of an ancient hybridization event that predates the differentiation of different river-basin specific populations of *S. pyrenaicus* and the species reaching its current distribution, as proposed by (Mendes et al., 2021; Mendes et al., in review).

Our results also suggest that the São Martinho population is an admixed population resulting from a much more recent hybridization event. This is supported by the parameter estimates of the demographic modeling, in agreement with the high levels of genetic diversity (Table S4), comparative lower levels of genetic differentiation (Table S3), as well as the PCA and ADMIXTURE analysis, where the São Martinho individuals show contribution from both *S. caetobrigus* and *S. pyrenaicus* Canha (ADMIXTURE analysis – Figures 2A, S4 and S11) or are placed in between the two species (PCA – Figures 2B, S5 and S9). Moreover, our D-statistic results indicate a similar contribution from both *S. caetobrigus* and *S. pyrenaicus* Canha into the genome of the São Martinho individuals (Figure 3B). While there is individual variation in parental contributions between São Martinho individuals, these contributions are more balanced and closer to 50/50 – for example, our estimates based on the ABBA/BABA patterns indicate the *S. pyrenaicus* contribution varies between 36% and 60% depending on the São Martinho individual analyzed (Figure 3D). We found no association between the mitochondrial type of each São Martinho individual (*S. caetobrigus* mitochondrial DNA or *S. pyrenaicus* mitochondrial DNA) and the contribution of each parental species to its nuclear genome (Figure 3D). However, we currently do not know how the ancestry tracts from each parental are distributed across the genome of the São Martinho individuals and how that distribution relates to the mitochondrial type of each individual.

Our demographic modelling consistently estimated similar parameter values across models for each hybridization case. For the *S. pyrenaicus* hybridization, our models consistently dated the divergence of *S. carolitertii* and *S. tartessicus* around 3 Mya. This is incongruent with previous works that estimated the divergence between these species to be around 6 Mya using seven nuclear genes (Sousa-Santos et al., 2019). However, the lower bound of the highest posterior density interval for such 6Mya estimate roughly matches our 3Mya estimates. Likewise, estimates of divergence between *S. pyrenaicus* and either of its sister species were estimated around 600,000-800,000 years ago, which closely fits the lower bound of the highest posterior density interval of previous studies (Sousa-Santos et al., 2019). The estimated contributions of *S. carolitertii* and *S. tartessicus* into the genome of *S. pyrenaicus* were also consistent across models and with the contribution estimated using the ABBA and BABA patterns but lower than those obtained using GBS data (Mendes et al., 2021). Our models never estimated more than 8% contribution from *S. tartessicus*, contrary to the findings of Mendes et al., 2021 which estimated 16-20% contribution from *S. tartessicus*. The majority of models also estimated a time of admixture around 600,000-800,000 years ago. This could be due to river captures that occurred between Tagus and both Guadiana and Douro that occurred in the past 500,000 years (Sousa-Santos et al., 2007). Surprisingly, ancient population sizes of *S. carolitertii* were constantly estimated to be higher than *S. tartessicus*. This was surprising as the genetic diversity of *S. tartessicus* is higher than *S. carolitertii* (Table S4). Likewise, Mendes et al., 2021 also estimated a larger ancestral size of *S. tartessicus*. However, we noticed that this estimate for *S. carolitertii* always approaches the maximum bound allowed for this estimate, which seems to imply that the ancestral population of *S. carolitertii* and ∼0.90 of *S. pyrenaicus* lineages was quite large. This indicates that some of the *S. pyrenaicus* lineages do not coalesce with *S. carolitertii* in the ancestral population, which could be an indication of a structured ancestral metapopulation or contribution from an unsampled ghost population related to *S. carolitertii*.

We observed that mostly our models considering admixture for the *S. pyrenaicus* hybridization case seemed to converge to one scenario, the Hybrid Origin model. We observed this with the secondary contact model between *S. carolitertii* and *S. pyrenaicus* (model Carol,(Pyr,Tar)+SecContCarol) and in both cases of divergence with gene flow between *S. carolitertii* and *S. tartessicus* (((Carol,Pyr),Tar)+SecContTar+AncMig); (Carol,(Pyr,Tar))+SecContCarol+AncMig). Not only do the times of admixture and divergence in these models overlap, but other parameters are mostly conserved between models (Table S9). This provided more confidence on the relationship between these three species, as independent models recovered the same topology. A recent work found similar model convergence in turbots and assumed this to reinforce the obtained relationship and estimates (Momigliano et al., 2024).

In the São Martinho hybridization, parameter estimates were also consistent between models. The divergence between *S. pyrenaicus* and *S. caetobrigus* was placed at 1 Mya. Previous works placed this divergence around the 6 Mya (Sousa-Santos et al., 2019), however, they also pose the possibility of migration between these two basins as during connectivity between the Lower Tagus paleobasin and the Alvalade paleobasin, or the Last Glacial Maximum due to regression of sea level or by connectivity established between Sado and left bank tributaries of Tagus. This could allow gene flow between these species, leading to more recent estimates of divergence. Furthermore, we observed that divergence with gene flow improved the likelihood of the models, especially due to gene flow from *S. pyrenaicus* into *S. caetobrigus*, with nearly no gene flow in the other direction. This suggests that *S. pyrenaicus* introgressed genetic material into *S. caetobrigus* but not the other way around. Since *S. caetobrigus* has a lower ancestral size, the lack of introgression into *S. pyrenaicus* may be due to selection purging introgressed alleles of *S. caetobrigus* into *S. pyrenaicus*. Admixture contribution from each parental was consistently estimated around 50% for both species (Figure 4), as happened in the contribution estimates using the ABBA and BABA patterns (Figure 3D) and the ADMIXTURE analysis (Figures 2 and S11). This also agrees with a more recent hybridization event, with more balanced contributions from both parentals. Corroborating this, our models date the hybridization event in the last 4,500 years (Figure 4 and S15). To our current knowledge, individuals undoubtably assigned to either one of the two parental species (*S. caetobrigus* and *S. pyrenaicus*) are not found in São Martinho. However, we cannot discard the hypothesis that 4,500 years ago, they did not coexist. Given our relatively low sample size, it is possible that individuals from both parental species are present in São Martinho, but due to their low frequency and density they have not been sampled. However, because our models suggest that hybridization happened around 4-5,00 years ago, we hypothesize that parental species are no longer present in São Martinho and instead have been replaced by individuals with admixed genomes. Balanced parental contributions into the genomes of the individuals of a hybrid lineage are often interpreted as a sign of very recent, possibly ongoing hybridization. However, if the pool of individuals at the initial hybridization event was comprised of *S. caetobrigus* and *S. pyrenaicus* in balanced numbers, the overall contribution of each species in the hybrid genomes may remain balanced even as the ancestry tracts become smaller in size due to recombination.

**Figure 4.**
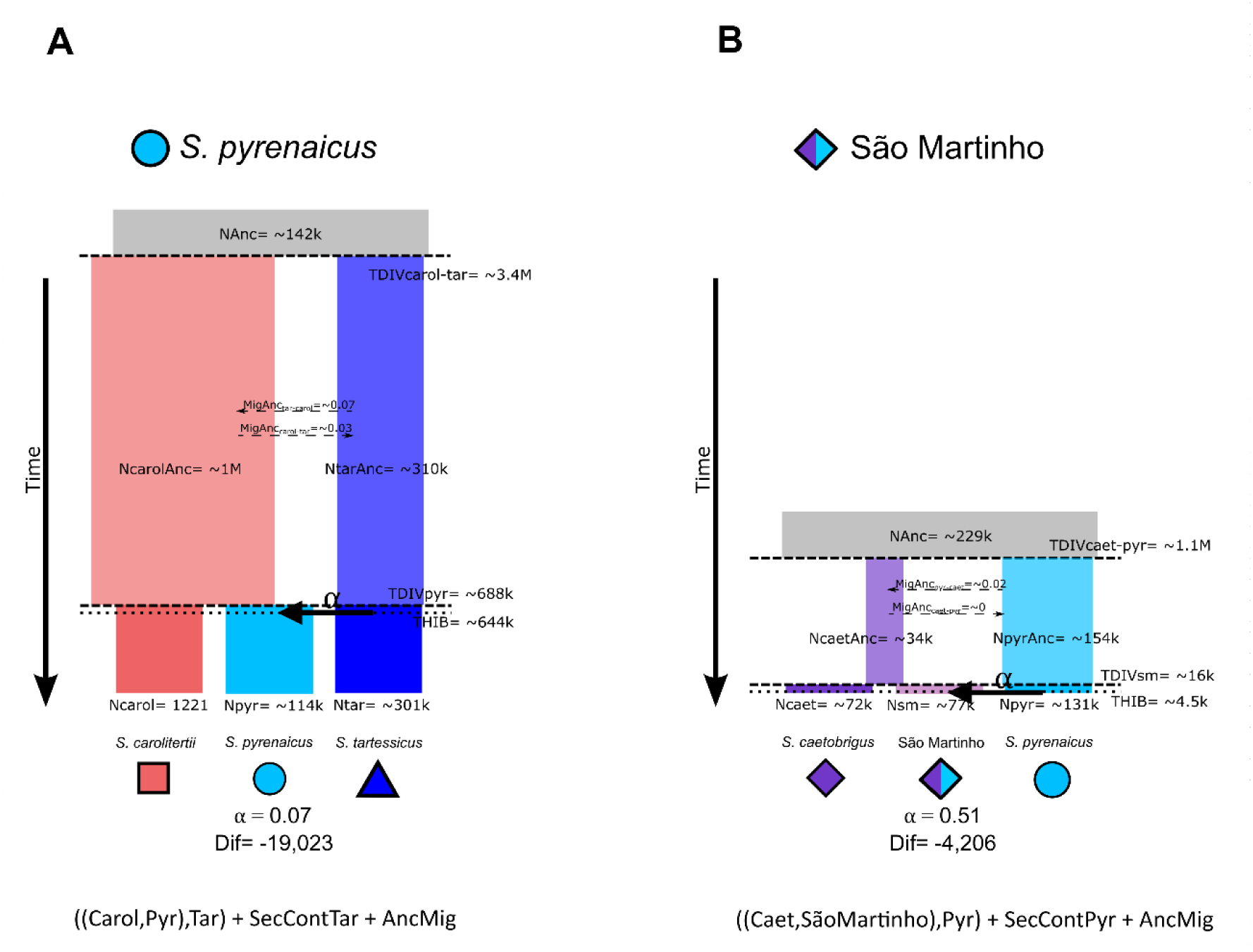
– Schematic representation of the likelihood parameters and parameters obtained of the best models tested with fastsimcoal2 for each hybridization case. (A) Best model obtained for the *S. pyrenaicus* hybridization case; (B) Best model obtained for the São Martinho hybridization case. The name given to each model is indicated below the model. For the results of all models tested for each hybridization case, see Figures S14 and S15 and Tables S9 and S10.

Our results might be affected by the assumptions of models and methods used, especially those of demographic modelling. First, there are currently no species-specific estimates of mutation rate for the Iberian chubs. Based on the estimates available of the literature for other fish species, we used a mutation rate of 1×10^-9^ in our demographic modelling. A recent study quantified germline mutation rates for eight species of Actinopterygii based on 19 trios (Bergeron et al., 2023), with an average per-generation mutation rate of 5.97 × 10^−9^ across these eight species. Available mutation rate estimates for sticklebacks (Zhang et al., 2023), Atlantic herring (Feng et al., 2017) and three species of Lake Malawi cichlids (Malinsky et al., 2018) also vary between 2×10^−9^ and 4.8×10^−9^. Thus, we believe our choice of mutation rate to be conservative within the range reported for other fish species. If a mutation rate for the Iberian chubs or a more closely related species becomes available, our estimates would need to be revised.

Similarly, to convert the divergence times into absolute time in Mya, we used a generation time of 3 years. This is commonly assumed to be a good estimate for the Iberian chubs and used as an average of all species. However, a precise estimation of the generation time in Iberian chubs, and whether this is the same for all species, is lacking. The information available suggests *S. carolitertii* males reach sexual maturity at two years, while females do so a year later, at 3 years (Alexandre et al., 2018; Maia, 2006). A similar pattern was found for *S. tartessicus*, where authors suggest fish from both sexes could mature in their second year of life, but the majority of fish and especially females did not mature until their third year (Pires et al., 2000). On the other hand, mean age at maturity was found to be 2 years for *S. torgalensis*, regardless of sex (Magalhães et al., 2003). Thus, the generation time used can also impact our estimates. Given the reported ages of maturity, it is likely that the generation time (average age of reproduction of females) is older, which would be reflected in older estimates of the time of events.

Furthermore, we used the genome of *Squalius cephalus*, the central European chub, as a reference to map our reads. The most recent estimates indicate the Iberian chubs lineage diverged from *S. cephalus* around 19Mya (Sousa-Santos et al., 2019). Our mapping statistics indicate around 2% mismatch between our study species and *S. cephalus* (Table S2). Currently, there is no available reference genome for any of the Iberian chub species. Thus, the *S. cephalus* reference genome is the closest available. This could potentially bias our estimates, and recent studies have looked into the effects of reference bias on various population genomics analyses (Prasad et al., 2022; Thorburn et al., 2023). We took steps to mitigate this as much as possible, namely by setting relatively high mapping quality thresholds and using the two closer outgroup species (*S. torgalensis* and *S. aradensis*) to ensure the correct polarization of the alleles.

Nevertheless, this could still have an impact on our estimates and, in the future, it would be interesting to see how our demographic modelling estimates would vary if a more closely related reference genome was used.

Finally, future work could be directed towards investigating the distribution of ancestry tracts from each parental species in the genomes of the two hybrid lineages we studied. Specifically, we currently do not know the size of these tracts or how they are distributed, for example, across genic and intergenic regions. Moreover, it would be interesting to investigate if there is repeatability both across individuals of these same hybrid lineage and between hybrid lineages.

## Acknowledgements

This work was funded by the Human Frontier Science Program Young Research Grant RGY0081/2020 and by the “Fundação para a Ciência e Tecnologia (FCT)” project HYBRIDOMICS (https://doi.org/10.54499/PTDC/BIA-EVL/4345/2021). This work was also funded through the FCT strategic projects UIDB/00329/2020 granted to cE3c (https://doi.org/10.54499/UIDB/00329/2020), UIDB/04292/2020 awarded to MARE (https://doi.org/10.54499/UIDB/04292/2020) and LA/P/0069/2020 granted to the Associate Laboratory ARNET (https://doi.org/10.54499/LA/P/0069/2020). We thank INCD (https://incd.pt/) for the use of their computing infrastructure, which FCT funded through project 2022.15858.CPCA.A2. SLM was funded by an individual FCT PhD scholarship (SFRH/BD/145153/2019).

